# Nest construction and its effect on post-hatching family life in the burying beetle *Nicrophorus vespilloides*

**DOI:** 10.1101/2024.03.29.587327

**Authors:** Eleanor Kate Bladon, Rebecca Mary Kilner

## Abstract

Through the effort required to construct them, the microenvironmental conditions they impose on the family and their indirect influence on post-hatching care, nests play a key role in influencing family life. We combined experimental evolution with cross-fostering experiments on laboratory populations of *Nicrophorus vespilloides* to investigate three ways in which the nest can contribute more broadly to parental investment. We used replicate populations of *N. vespilloides* that had evolved for 42 generations under contrasting regimes of care. Populations were either able to supply post-hatching care (“Full Care”) or prevented from supplying any post-hatching care (“No Care”). Research on these populations has previously shown that the No Care populations evolved to build rounder nests, more rapidly, by Generation 14. Here we found: 1) larvae raised by Full Care parents on nests prepared by parents from the No Care population did not attain a higher mass by the end of larval development than larvae in other treatments. However, we did discover that: 2) cross-fostering nests between families consistently reduced larval mass – and to a similar extent whether nests were cross-fostered between or within the populations. We suggest that cross-fostering disrupted the chemical environment on and around the nest since we found no evidence that 3) nests mediate interactions between males and females. The duration of paternal care was consistently shorter than the duration of maternal care, and even shorter for males from the No Care populations than males from the Full Care populations. Nevertheless, the duration of male care did not predict variation in duration of female care. In short, although the nest is the substrate for burying beetle family life, we found little evidence that it had evolved divergently in our experimental populations to influence parental investment.

## Introduction

Across taxa, nests play a central role in family life by providing a nursery in which young animals develop (Soler et al., 1998). Although they vary greatly in their complexity, nests are a key component of pre-hatching parental investment, and the key site for post-hatching parental investment, mediated by social interactions within the family. Through the effort required to construct them, the microenvironmental conditions they impose on the family and their indirect influence on post-hatching care, nests potentially play a key role in regulating family life (Hansell, 2005). Yet these qualities are typically tightly correlated with each other and with parental quality. Teasing apart the separate and diverse effects of nest construction on animal family life therefore requires an experimental approach. In this paper we combine experimental evolution with cross-fostering experiments to investigate the extent to which the nest contributes to parental investment more broadly.

The first question we consider centres on the role that nests play in mediating pre- and post-hatching contributions to offspring fitness. Nest construction is a type of pre-hatching parental investment since it promotes offspring fitness but at a fitness cost to the parents (Clutton-Brock, 1991; De Gasperin et al., 2016). Nests promote fitness because they buffer vulnerable offspring from predation, intraspecific competition and environmental stressors (Hilton et al., 2004; Magee & Neff, 2006; Wood & Bjorndal, 2000). Nest construction is costly for adults, both energetically (Benowitz et al., 2017; De Gasperin et al., 2016; Mainwaring & Hartley, 2013) and because adults sustain an increased the risk of attack whilst nest-building (e.g. Berger-Tal et al., 2010; Chalfoun & Martin, 2010). Just as with other forms of pre-hatching investment (Krist, 2011; Mäenpää & Smiseth, 2017; Paquet & Smiseth, 2016), investment in nest building must be balanced against the supply of post-hatching parental investment. Here we investigate how the two forms of investment combine to influence offspring fitness.

The effects of nest construction can endure to influence life after hatching too. Nests provide the physical environment inhabited by families as their offspring develop, and are therefore a source of selection on the family during this sensitive period. Across bird species, the nestlings’ begging calls and visual displays are correlated with the nest environment - the calls being louder when the nest is safer from predators (Briskie et al., 1999), and mouthparts showing greater brightness contrast when the nest interior is poorly lit (Kilner & Davies, 1998). Within bird species, individuals have been shown to co-adapt their behaviour to the social environment within their own family (Royle et al., 2012). Here we investigate whether families within species are similarly co-adapted to the nest they have constructed for themselves.

Nests have also been shown to mediate social interactions within the family through the structural barriers they introduce that constrain, or reinforce, power asymmetries when food is distributed among offspring. Bird nest architectures create single points of food delivery, for example, which can be monopolised by competitively dominant offspring (Kacelnik et al., 1995; Ostreiher, 1997) but which can also provide different feeding zones for each parent, enabling sex-specific provisioning rules to operate (Tanner et al., 2008). The discrete cells that make up social insect nests, on the other hand, limit the extent of competition within the brood. Here we take a different perspective by considering whether the burying beetle’s edible nest can contribute to post-hatching interactions within the family, because it is the source of nourishment supplied during post-hatching care.

We describe an experiment on the burying beetle *Nicrophorus vespilloides* that addresses each of these three issues. To reproduce, burying beetles locate a small dead vertebrate and then convert the corpse into an edible nest for their larvae. Together, a pair will rip off its fur or feathers, spread the flesh with antimicrobial oral and anal fluids, roll it into a ball and bury it underground (Cotter & Kilner, 2010; Scott, 1998). During this process they mate, and the female lays her eggs in the soil around the prepared carcass nest. When they hatch, the larvae crawl to the carcass where they feed themselves on the flesh and parents feed them trophallactically (Eggert et al., 1998). There is some sexual division of care within pairs, with males performing more of the carcass preparation and larval defence activities, and females providing more of the direct larval feeding (Royle & Hopwood, 2017; Smiseth et al., 2006; Walling et al., 2008). Larval development is complete about a week after hatching, at which point larvae start to crawl away to pupate in the soil nearby and parents fly off in search of new reproductive opportunities.

This suite of behaviours is believed to be the most common in nature, but variations also arise, such that larvae may receive post-hatching care from only one parent, or no post-hatching care at all (Müller et al., 2007). If a female with stored sperm finds a carcass with no males present, she can prepare the nest herself, lay fertilised eggs, and provide uniparental care (Eggert & Müller, 1997). Both males and females may also end up providing uniparental care if one of them dies or deserts before the eggs hatch. We exploited this variation in parental care to establish replicate evolving populations in the laboratory, in which we manipulated the supply of care. In two populations (“No Care” populations), parents were prevented from providing any post-hatching care, whereas in two other populations (“Full Care” populations), parents were allowed to stay with their offspring throughout their development. After 14 generations of experimental evolution, Duarte et al. (2021) found that No Care parents had evolved to prepare the carcass nest more rapidly than Full Care parents. However, by generation 43, relaxed selection had caused parental care behaviours to decay in No Care fathers (Bladon et al., 2023). The knock-on consequences of these changes in carrion nest preparation, combined with changes in post-hatching care, for interactions within the family have not yet been explored. We investigated these effects by swapping carcass nests within and between experimental populations, after 41 generations of experimental evolution. We then asked:

### Pre-hatching care and offspring performance

How does evolved change in pre-hatching care in the No Care populations combine with post-hatching care to influence offspring performance? If more effective pre-hatching care combines additively with post-hatching care, then we would expect broods to attain the greatest mass when they receive post-hatching care from Full Care parents whilst occupying a No Care-prepared carcass, all else being equal.

#### Are families co-adapted to their carrion nest?

We hypothesise that families co-adapt to the carrion nest environment they construct. Co-adaptation might be so subtly attuned that it varies from one family to the next. Any differences seen are likely to be mediated by the fluids that the parents spread on the carcass during and after nest preparation, as well as any chemical trails they leave in the surrounding soil to guide their newly hatched larvae to the carrion nest (Fouche et al., 2018; Gruszka et al., 2020). Previous research in this and other *Nicrophorus* species has shown that the fluids change the chemical signature emanating from the carcass (Trumbo et al., 2021) and the microbial community inhabiting it (Wang & Rozen, 2017). The fluids not only suppress competitive bacteria, but are also an indirect vertical method of transmission of beneficial symbionts and proteins to the larvae (Jacobs et al., 2016; Körner et al., 2023; Shukla et al., 2017). An experiment by Arce et al. (2012) showed that larvae reared without parental care on liver pieces spread with either beetle anal secretions, hen egg-white lysozyme or PBS had the highest survival in the anal secretion treatment, indicating that it is not the bacterial-killing properties of the exudates’ lysozyme activity alone that increase larval fitness. To our knowledge no studies have tested the effects on larval fitness of giving larvae carcasses spread with the fluids of adults that are not their parents, compared to carcasses spread with their parents’ fluids. Since the fluids on the carcass represent an important method of beneficial microbe transmission, it is possible that larvae are co-adapted to the specific cocktail of microbes or chemicals in these fluids produced by their parents. If this is true, then we predict that swapping nests between families, whether they are from the same population or not, will cause broods to be smaller and lighter at dispersal. Alternatively, co-adaptation might only be detectable at the population level. If this is true, then we predict that swapping nests between No Care and Full Care populations will cause broods to be lighter at dispersal, regardless of whether the nests were originally prepared by the No Care or Full Care populations.

#### Does the carrion nest mediate interactions between the parents?

Across species that show biparental care, the care provided by one parent can influence the care provided by the other through the phenomena of behavioural ‘matching’ and ‘negotiation’ (Paquet & Smiseth, 2016). Indeed, previous work on burying beetles suggests that investment in care by one parent can influence the care provided by the other (Cotter & Kilner, 2010). Therefore, we predicted that the duration of care supplied by each parent should be influenced by the effort of its partner, but that this relationship could be lost when nests are swapped between families if parental behavioural synchronicity post-hatching is mediated by chemical cues laid down in the parents’ oral fluids during nest preparation. We further predicted that parents from the No Care populations would be less sensitive to each other, and to the carrion nest, considering that the male’s capacity to supply post-hatching care has decayed, while the female’s has not (Bladon et al., 2023).

## Methods

### Experimental Evolution

The *N. vespilloides* populations described in this study were part of a long-term experimental evolution project that investigated how populations of burying beetles adapt to the loss of post-hatching parental care. It focused on four experimental populations: Full Care (FC, x2 replicates) and No Care (NC, x2 replicates). Their establishment and husbandry have been described in detail before (Duarte et al., 2021; Jarrett, Evans, et al., 2018; Jarrett, Rebar, et al., 2018; Rebar et al., 2020; Schrader, Cosby, et al., 2015; Schrader, Jarrett, et al., 2015b; Schrader et al., 2017). Briefly, these populations were established in 2014 with wild-caught beetles (trapped under permit) from four woodland sites across Cambridgeshire, UK (Byron’s Pool, Gamlingay Woods, Waresley Woods and Overhall Grove). The No Care populations were routinely prevented from supplying any post-hatching care, through the removal of adults at 53h post-pairing, when the carrion nest was complete but before the larvae had hatched, whereas in the Full Care populations adults were allowed to stay with their larvae throughout development and provide care. This procedure was repeated at every generation. Each type of experimental population was run in a separate block (FC1/NC1 and FC2/NC2) with breeding staggered between blocks by 7 days.

### General husbandry

For maintaining the populations at each generation, pairs of unrelated sexually mature male and female beetles from the Full Care and No Care populations were bred by placing them in plastic breeding boxes (17 x 12 x 6 cm) with damp soil (Miracle-Gro Compost) and 10-15 g mouse carcasses on which to breed. Parents were removed from the No Care boxes at 53 h as described above. At natural dispersal time (8 days after pairing), larvae were counted, weighed and placed in plastic pupation boxes (10 x 10 x 2 cm), filled with damp peat. Sexually immature adults were eclosed approximately 21 days later and housed individually in separate boxes (12 x 8 x 2 cm). Adults were fed twice a week (beef mince) until breeding, which took place 15 days post-eclosion. Adults and pupating larvae were kept on a 16L: 8D hour light cycle at 21°C.

### Experimental design

Experimental beetles for this study were derived from generation 42 of the experimental evolution populations. Sexually mature pairs of No Care or Full Care beetles were placed with a 9-12 g mouse carcass in a large breeding box with a plastic partition and one-way valve that allowed parents to desert the nest after they ceased parental care, but did not allow them to return (De Gasperin & Kilner, 2015; Figure A1). The mouse was weighed before it was added to the box.

At 53 h post-pairing, some prepared carcasses were transferred between boxes, while the clutch remained in the soil, in the box with the biological parents. This meant that some pairs and eggs kept their own prepared carcass, some received a carcass prepared by a pair from the same experimentally evolving population, and some received a carcass prepared by a pair from the other experimentally evolving population (see Figure A2 for overview of experimental design). Those carcasses that were not transferred were lifted out of the box and immediately returned to ensure the same level of handling across all broods. Following the transfer, adults remained in the boxes until they entered the escape chamber or the experiment ended (8 days after pairing). Thus, in all treatments parents had the opportunity to supply post-hatching care. From breeding to dispersal, the escape chamber was checked every 4 hours between 8am and 8pm (i.e. 56-192 h after pairing). Their time of first detection in the escape chamber was used to infer the duration of parental care (De Gasperin & Kilner, 2015). Eight days post-pairing, remaining parents were removed, the number of surviving larvae was counted and the brood weighed. This design enabled us to partition the contributions of pre-hatching and post-hatching components of parental investment to brood mass and size at dispersal. Brood mass is a key correlate of fitness because larval mass increases with the supply of parental care (Bladon et al., 2020; Pascoal et al., 2018; Steiger, 2013) and larvae that are larger at dispersal eclose as larger adults (Jarrett et al., 2017) with higher fecundity (Bladon et al., 2020; Pascoal et al., 2018).

### Ethical note

The beetles that established the experimental populations were caught in carrion-baited traps under permit from Natural England and with landowner permission. Larvae and beetles in the experimental populations, and in the subset used for experiments, were handled carefully and as minimally as possible and had access to plenty of food. At all life stages, beetles were kept in containers which had sufficient space and had been thoroughly cleaned prior to housing them. During the breeding experiment the maximum length of time that a beetle could have been in the escape chamber without food would have been 12 h. At the end of the breeding experiment, beetles and larvae were placed in a -20 °C freezer to euthanise them. Mice used for population maintenance and experiments were bought from a commercial reptile food supplier.

### Statistical analyses

All statistical tests were conducted in R version 4.2.0 (R Core Team, 2022). Data handling and visualisation were carried out using base R and the ‘tidyverse’ suite of R packages (Wickham et al., 2019). Model selection for the analyses of brood mass and brood size was conducted using the ‘dredge’ function in the ‘MuMIn’ R package (Bartoń, 2023). The most parsimonious model within two AIC_C_ points of the model with the lowest AIC_C_ was chosen as the optimal model. Goodness of fit of optimal models was assessed with the ‘DHARMa’ R package (Hartig, 2022). In the survival models analysing leaving time, a stepwise deletion method using *F* tests, implemented in the base ‘statistics’ package in R, was used to determine significance of each term and remove non-significant terms sequentially (Crawley, 2007). Continuous independent variables in all models were scaled and centred using the ‘scale’ function in base R. Collinearity was assessed using the ‘vif’ function in the ‘car’ package, and the highest variance inflation factor for any variable in a model was 1.175, indicating no issues with multicollinearity. The biological reason for including each variable in the model is given in Table A1.

### Pre-hatching care and offspring performance

#### Are families co-adapted to their carrion nest?

To address these, brood mass and brood size were analysed using linear models with Gaussian error structures. In separate models, either brood mass (mass of all larvae) or brood size (number of larvae) at dispersal was used as the dependent variable, and the independent variables included in the maximal model were original carcass mass (g) (i.e. the mass of the mouse given to parents at the start of the experiment), current carcass mass (g) (i.e. the original mass of the mouse that has become the nest currently used by the focal parents), parents’ experimental population of origin (No Care or Full Care), carcass preparers’ experimental population of origin (No Care or Full Care), whether the carcass had been transferred between broods or not, male leaving time, female leaving time and experimental block (1 or 2). We also included the interactions between: parents’ experimental population and whether the carcass had been transferred, the masses of the original and current carcasses, the parents’ experimental population and male leaving time parents’ experimental population and female leaving time, and parents’ experimental population and carcass preparers’ experimental population. In the model of brood mass, brood size was also included as an independent variable.

#### Does the carrion nest mediate interactions between the parents?

A t-test was implemented (using the base R ‘statistics’ package) to determine whether there was a significant difference in the duration of male and female care. As there was a significant difference, male and female leaving times were analysed separately in subsequent analyses. Survival models were used for analysing the duration of care. Hazards did not conform to any parametric distribution, but their proportionality was checked using the cox.zph function in the survival R package (Therneau, 2022) and by generating scaled Schoenfeld residuals (Therneau & Grambsch, 2000), indicating that for both males and females the hazards were proportional to the independent variables. Therefore, data were analysed using semi parametric Cox’s proportional models for interval censored data, using the ‘icenReg’ R package (Anderson-Bergman, 2017). In both maximal models, the predictor variables were focal individual’s (male or female) experimental population (No Care or Full Care), carcass preparers’ experimental population (No Care or Full Care), brood size, original carcass mass (g), current carcass mass (g), whether the carcass had been transferred, brood size, experimental block (1 or 2), as well as interactions between: the focal individual’s experimental population and whether the carcass had been transferred, the mass of the original and current carcass, and the focal individual’s experimental population and carcass preparers’ experimental population. Most males left before females (*N* = 202/218 pairs), therefore female duration of care could not be included in the male care duration model, but male duration of care was included in the maximal female care duration model as an independent variable, along with the interactions between male experimental population and his leaving time, and male leaving time and whether the carcass was transferred.

## Results

### Pre-hatching care and offspring performance

We found no evidence that carcasses prepared by No Care parents compensated fully for worse post-hatching care. The optimal model of brood mass retained the original carcass mass, current carcass mass, parents’ experimental population, whether the carcass was transferred, female leaving time and brood size as independent variables (Tables 1 & A2). No Care larvae were lighter than Full Care larvae, and there was no increase in larval mass in either population type when they developed on a carcass prepared by No Care parents (Figure 1).

**Figure 1:**
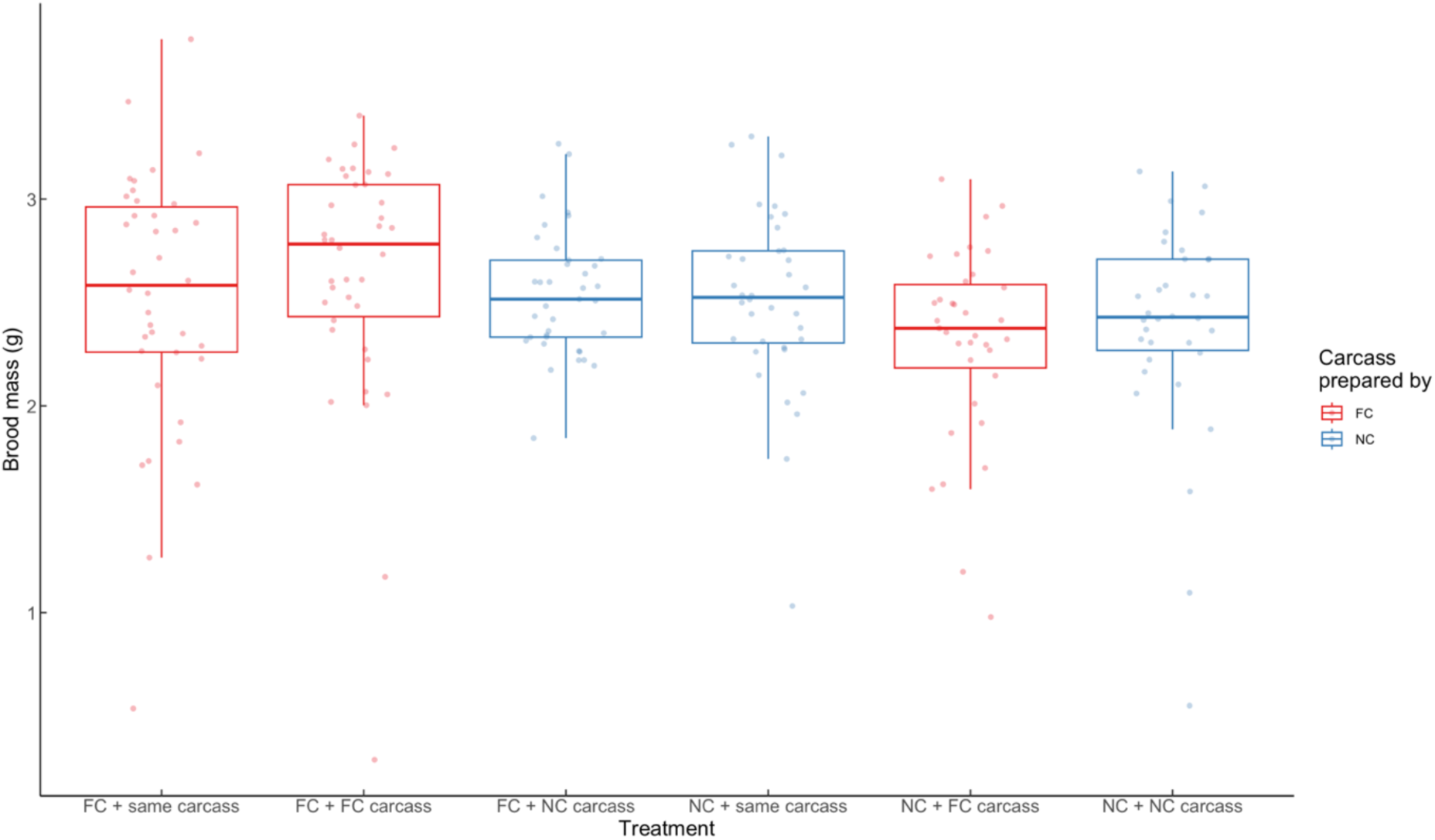
The effect of carcass translocation treatment on brood mass at larval dispersal. For treatments shown in red, carcasses were prepared by pairs from the Full Care lines. For treatments shown in blue, carcasses were prepared by pairs from the No Care lines. In “FC + same carcass” and “NC + same carcass” treatments the carcasses remained with the pairs that had prepared them. In the other treatments, focal pairs received a nest prepared by a different pair, either from the same or from a different experimental population as themselves. Whiskers extend to the farthest data point which is no more than 1.5 times the interquartile range from the box. Points represent data for individual broods. *N* = 218 broods.

**Table 1:** Results of a Gaussian linear model analysing predictors of brood mass at dispersal. Terms retained in the optimal model are presented. All continuous variables were scaled and centred.

| Independent variable | Estimate | SE | *p* |
| --- | --- | --- | --- |
| Intercept | 2.599 | 0.032 | <0.001 |
| Original carcass mass | 0.036 | 0.017 | 0.034 |
| Current carcass mass | 0.076 | 0.017 | <0.001 |
| Parents' experimental population (NC) | -0.128 | 0.032 | <0.001 |
| Carcass transferred? (Yes) | -0.074 | 0.034 | 0.028 |
| Female leaving time | 0.003 | 0.001 | 0.013 |
| Brood size | 0.073 | 0.003 | <0.001 |

There was a positive relationship between the mass of the current carcass and brood mass (Figure 2a). Furthermore, when pairs prepared a larger carcass their broods were subsequently heavier (Figure 2b), even if that carcass was transferred away from them before the larvae hatched.

**Figure 2:**
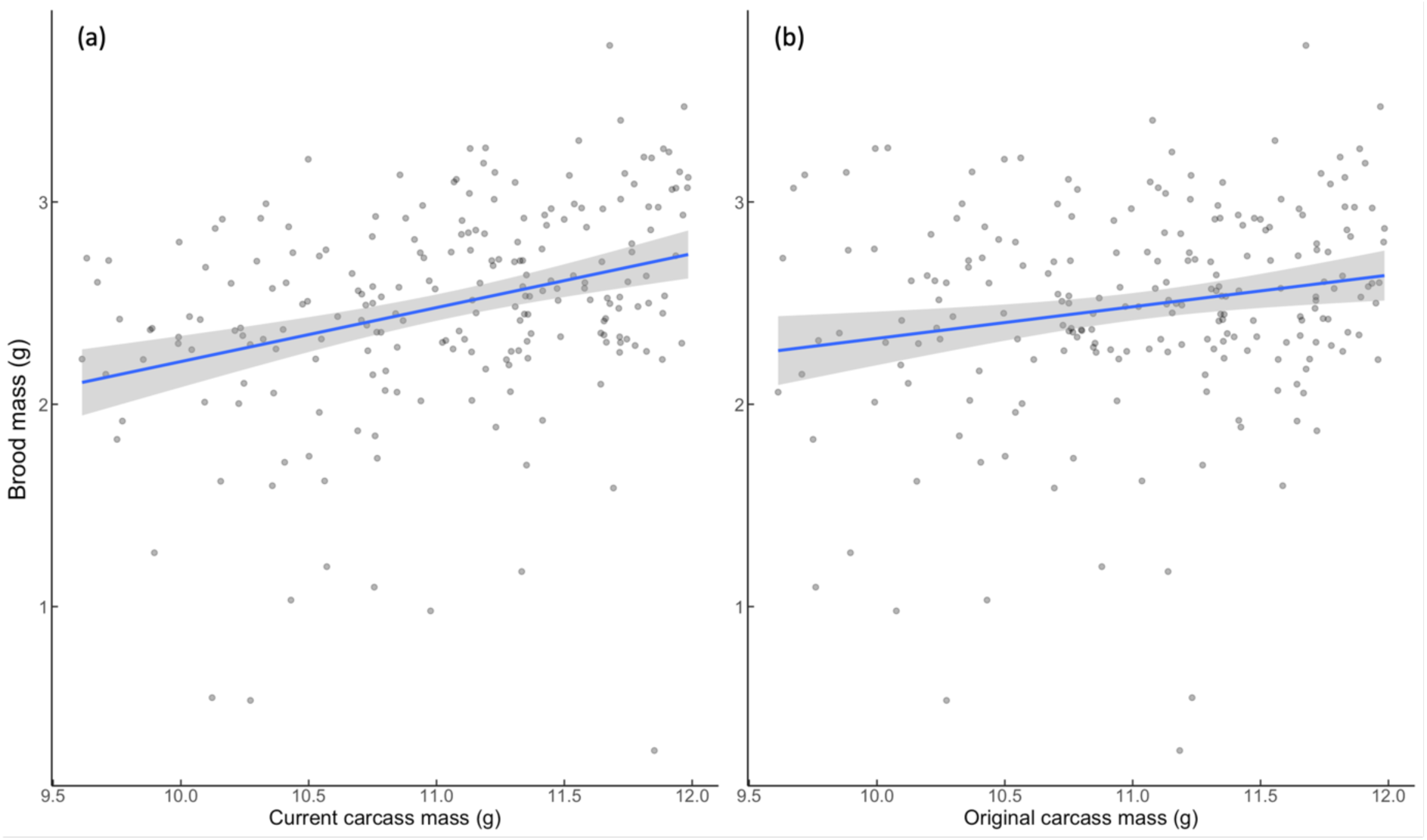
The effect of a) the mass of the carrion nest used by focal parents during post-hatching care (measured when it was a mouse, at the start of the experiment) and b) the mass of the mouse before it was converted into a carrion nest by focal parents. Shaded areas indicate the 95 % confidence intervals. In all plots points represent data for individual broods. *N* = 218 broods.

The optimal model of brood size retained current carcass mass, male leaving time and experimental block as independent variables (Tables 2 & A3). As expected, the linear model analysing brood size indicated that current carcass mass was positively correlated with brood size. However, contrary to predictions, experimental population (Full Care or No Care), of either the caring parents or the carcass preparers, was not retained in the optimal model, either as a main effect or in an interaction (Figure 3).

**Figure 3:**
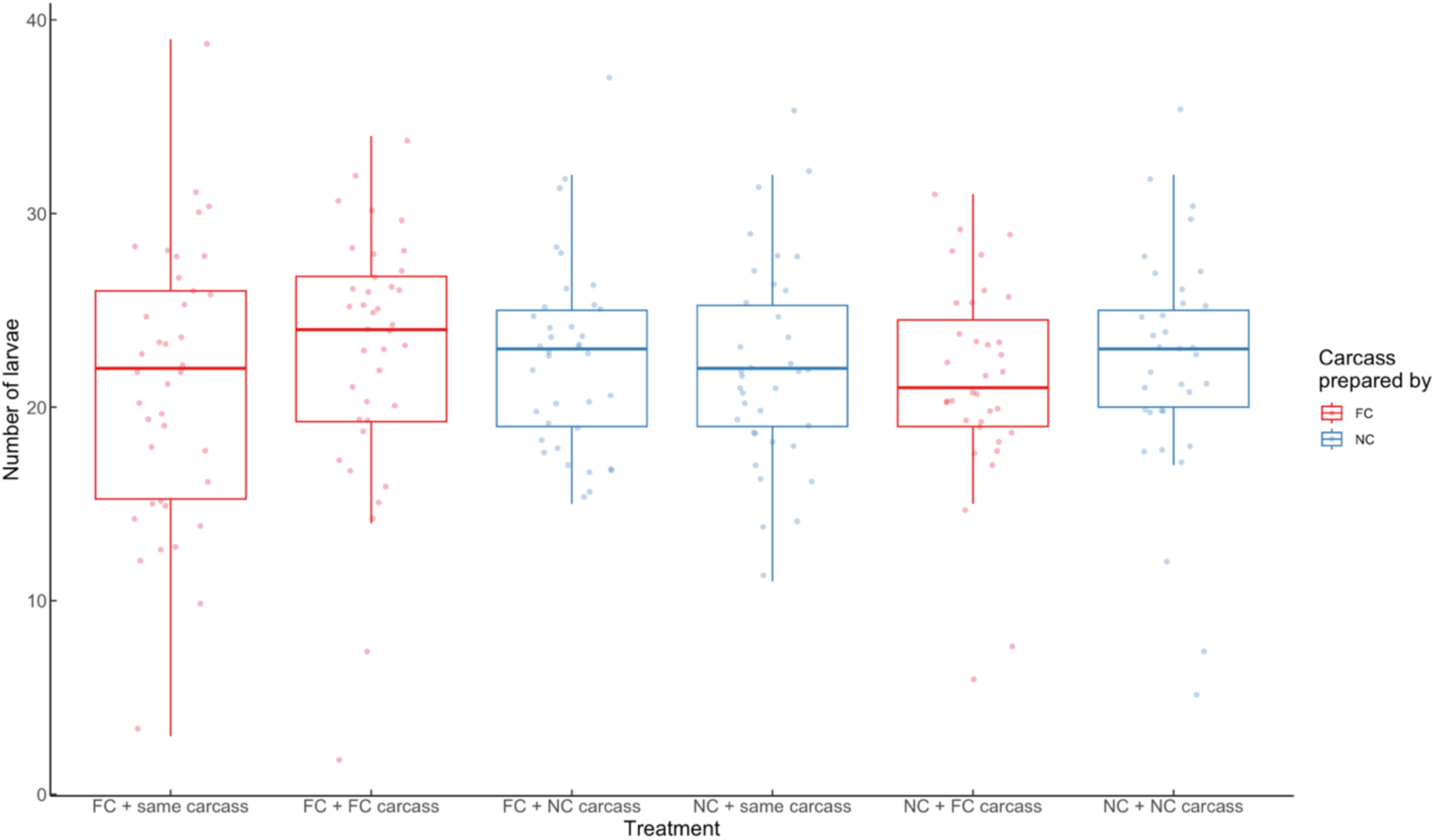
The effect of carcass translocation treatment on brood size at dispersal. For treatments shown in red, carcasses were prepared by pairs from the Full Care lines. For treatments shown in blue, carcasses were prepared by pairs from the No Care lines. In “FC + same carcass” and “NC + same carcass” treatments the carcasses remained with the pairs that prepared them. In the other treatments, focal pairs received a carcass that had been converted into a nest by a different pair, which was either from the same or from a different experimental population as themselves. Whiskers extend to the farthest data point which is no more than 1.5 times the interquartile range from the box. Points represent data for individual broods. *N* = 218 broods.

**Table 2:**
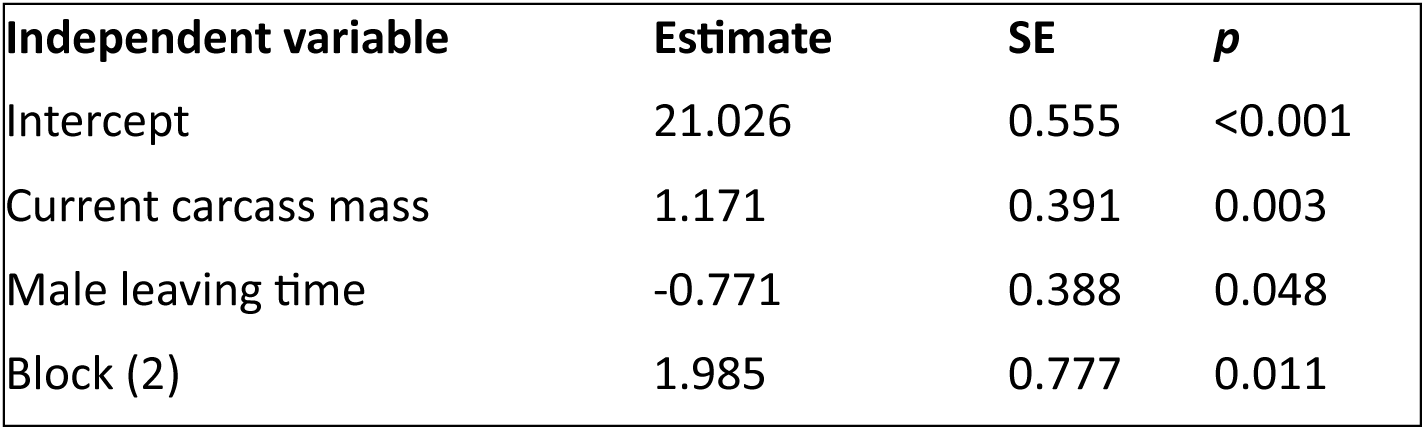
Results of a Gaussian linear model analysing predictors of brood size at dispersal. Terms retained in the optimal model are presented. All continuous variables were scaled and centred.

| Independent variable | Estimate | SE | $p$ |
| --- | --- | --- | --- |
| Intercept | 21.026 | 0.555 | <0.001 |
| Current carcass mass | 1.171 | 0.391 | 0.003 |
| Male leaving time | -0.771 | 0.388 | 0.048 |
| Block (2) | 1.985 | 0.777 | 0.011 |

#### Are families co-adapted to their carrion nest?

When pairs from either type of population bred on a carcass that had been transferred to them (rather than on the carrion nest that they themselves had prepared) they produced lighter broods at dispersal (Figure 4) regardless of their population of origin. There was no evidence that the extent of co-adaptation differed between the different types of experimental population. The interaction between experimental population and whether the carcass had been transferred was not retained (Tables 1 & A2).

**Figure 4:**
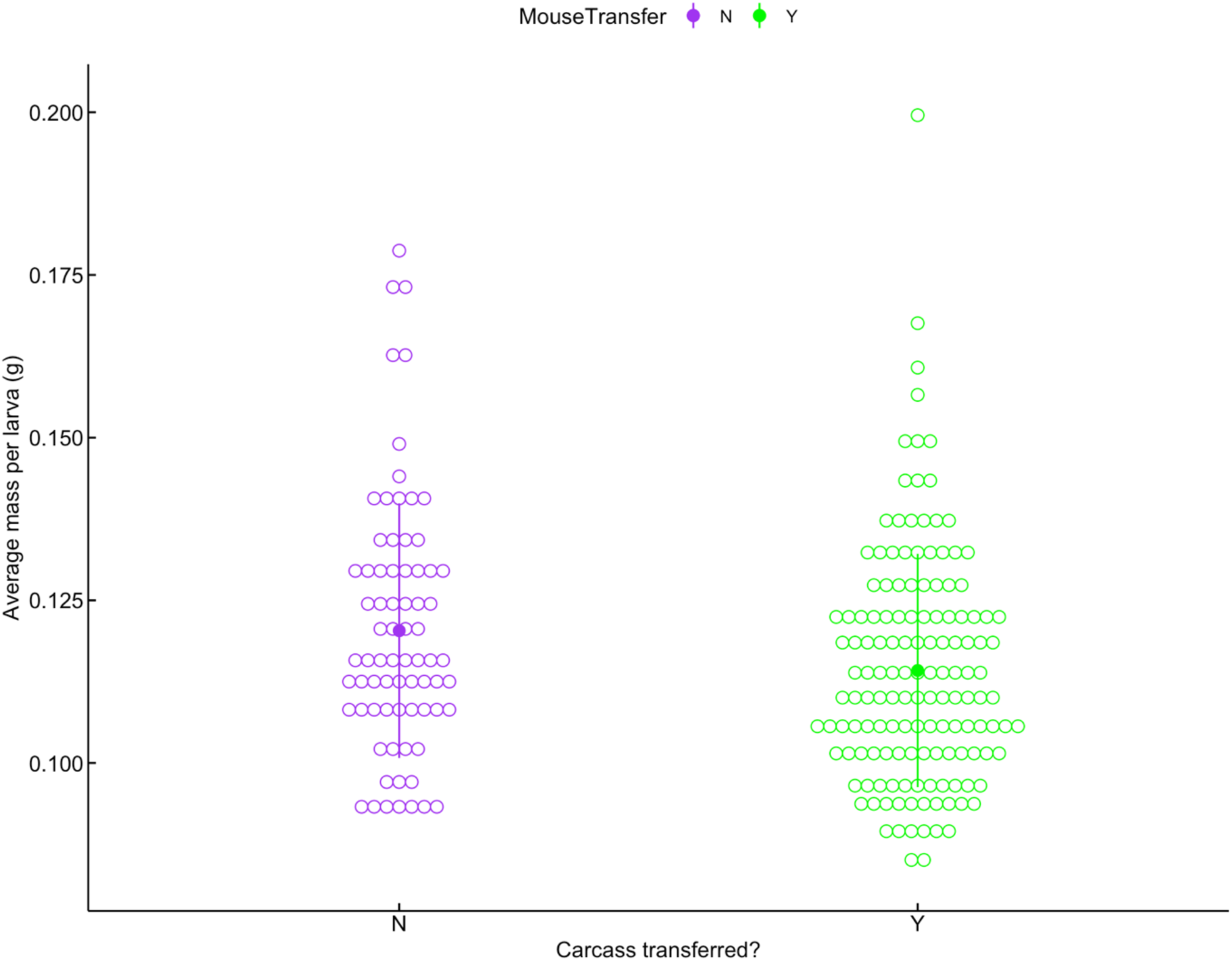
The effect of whether the carcass was translocated on the average mass of each larva at dispersal, derived from brood mass/brood size. Open circles represent raw data points for each brood, filled points represent the mean for each treatment and vertical lines represent the standard deviation. Purple = carcass not transferred, green = carcass transferred. *N* = 218 broods.

#### Does the carrion nest mediate interactions between the parents?

In general, and regardless of their experimental population, males left their brood earlier (M = 144, SD = 35.9) than females (M = 184, SD = 15.2). The difference, -40.032 hours (95% CI [-45.143, -34.921]), was statistically significant; t(217) = -15.436, *P* = <0.001.

Male departure time was predicted by an interaction between the experimental population of the resident male on the carcass and the experimental population of the pair that prepared the carcass (Figure 5). On carcasses prepared by Full Care pairs, Full care males stayed longer than No Care males, but on carcasses prepared by No Care pairs both No Care and Full Care males stayed for similar lengths of time (interaction term: hazard ratio = 0.516, Wald = -2.218, *P* = 0.027). The same result was not found for females (Table 3).

**Figure 5:**
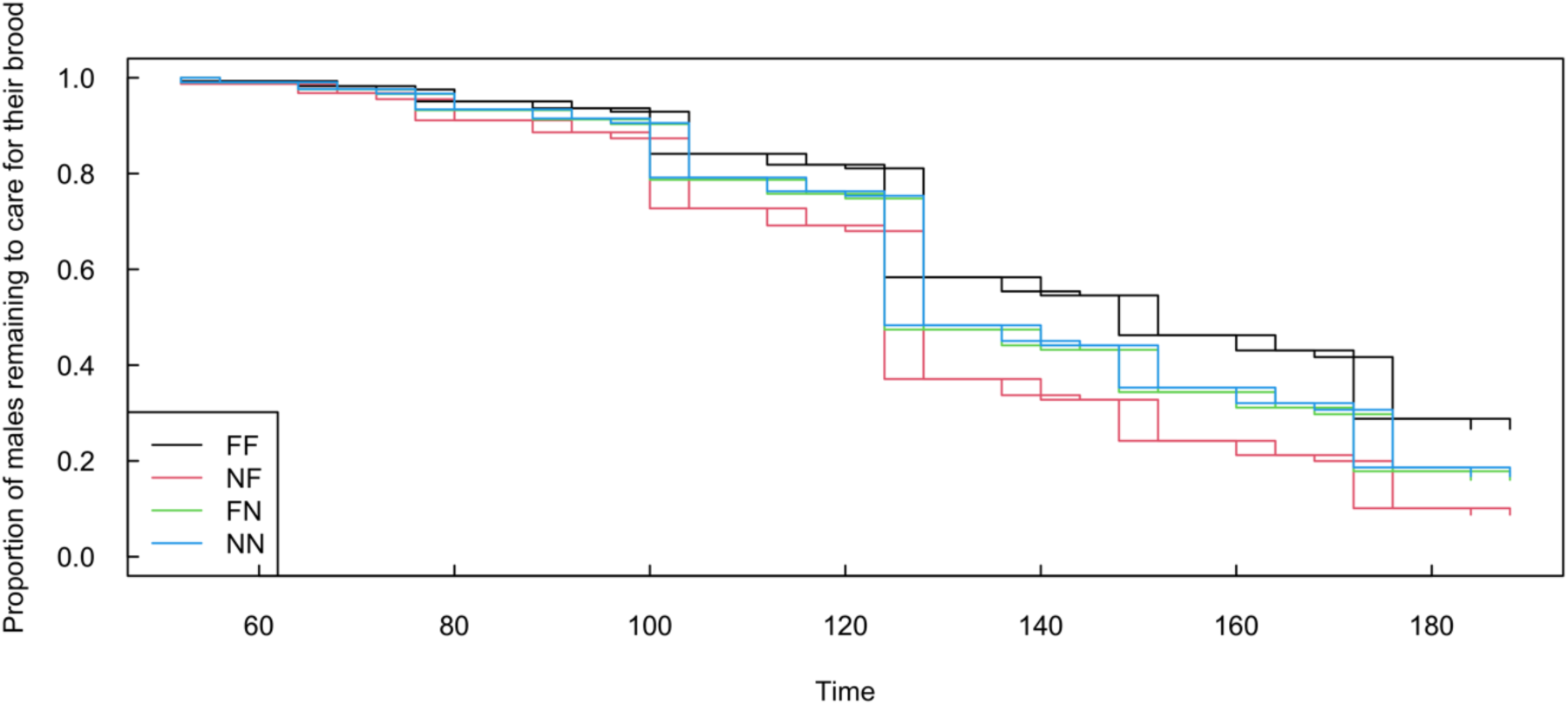
The effect of the interaction between the experimental population of the carcass preparer and the male resident on the duration of his care. FF: nest prepared by Full Care parents, then used by Full Care parents to raise larvae; NF: nest prepared by Full Care parents, then used by No Care parents to raise larvae; FN: nest prepared by No Care parents, then used by Full Care parents to raise larvae; NN nest prepared by No Care parents, then used by No Care parents to raise larvae. Two lines are displayed for each treatment because with interval censored data it was not possible to tell the exact time that the male left, only the time period within which he left. The lines depicted are predicted by the model fitted to the data: for any point between the two lines there is the same likelihood of the parent remaining. *N* = 218 broods.

**Table 3:**
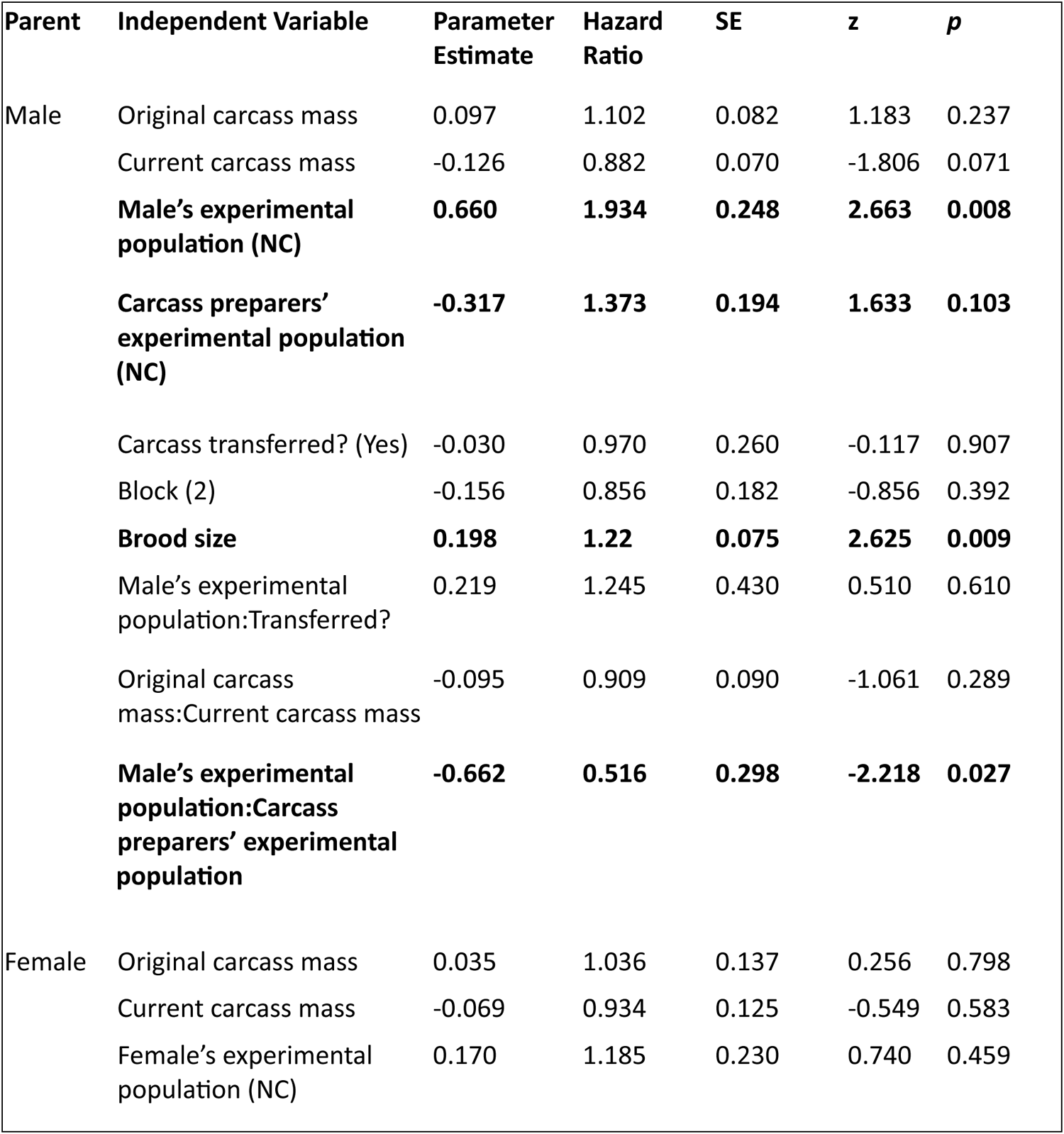
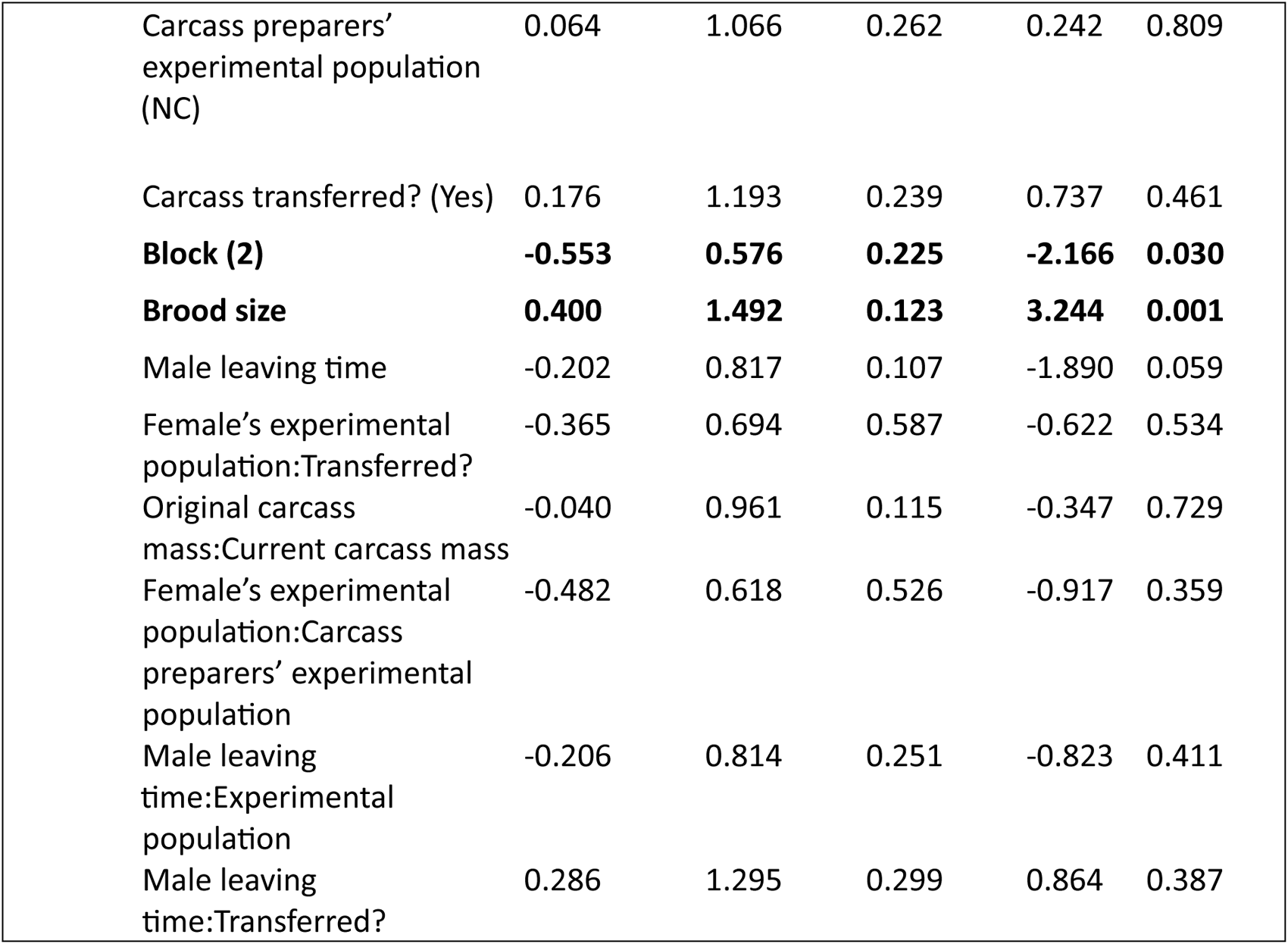
Results of semi-parametric Cox’s proportional models of parental leaving times. Terms retained in the minimal model are shown in bold. All terms included in the maximal model are given and statistics provided are for the last model in which the term was retained “:” represents an interaction between terms. All continuous variables were scaled and centred.

| Parent | Independent Variable | Parameter Estimate | Hazard Ratio | SE | z | p |
| --- | --- | --- | --- | --- | --- | --- |
| Male | Original carcass mass | 0.097 | 1.102 | 0.082 | 1.183 | 0.237 |
|  | Current carcass mass | -0.126 | 0.882 | 0.070 | -1.806 | 0.071 |
|  | **Male's experimental population (NC)** | **0.660** | **1.934** | **0.248** | **2.663** | **0.008** |
|  | **Carcass preparers' experimental population (NC)** | **-0.317** | **1.373** | **0.194** | **1.633** | **0.103** |
|  | Carcass transferred? (Yes) | -0.030 | 0.970 | 0.260 | -0.117 | 0.907 |
|  | Block (2) | -0.156 | 0.856 | 0.182 | -0.856 | 0.392 |
|  | **Brood size** | **0.198** | **1.22** | **0.075** | **2.625** | **0.009** |
|  | Male's experimental population:Transferred? | 0.219 | 1.245 | 0.430 | 0.510 | 0.610 |
|  | Original carcass mass:Current carcass mass | -0.095 | 0.909 | 0.090 | -1.061 | 0.289 |
|  | **Male's experimental population:Carcass preparers' experimental population** | **-0.662** | **0.516** | **0.298** | **-2.218** | **0.027** |
| Female | Original carcass mass | 0.035 | 1.036 | 0.137 | 0.256 | 0.798 |
|  | Current carcass mass | -0.069 | 0.934 | 0.125 | -0.549 | 0.583 |
|  | Female's experimental population (NC) | 0.170 | 1.185 | 0.230 | 0.740 | 0.459 |

|  |  |  |  |  |  |
| --- | --- | --- | --- | --- | --- |
| Carcass preparers' experimental population (NC) | 0.064 | 1.066 | 0.262 | 0.242 | 0.809 |
| Carcass transferred? (Yes) | 0.176 | 1.193 | 0.239 | 0.737 | 0.461 |
| **Block (2)** | **-0.553** | **0.576** | **0.225** | **-2.166** | **0.030** |
| **Brood size** | **0.400** | **1.492** | **0.123** | **3.244** | **0.001** |
| Male leaving time | -0.202 | 0.817 | 0.107 | -1.890 | 0.059 |
| Female's experimental population:Transferred? | -0.365 | 0.694 | 0.587 | -0.622 | 0.534 |
| Original carcass mass:Current carcass mass | -0.040 | 0.961 | 0.115 | -0.347 | 0.729 |
| Female's experimental population:Carcass preparers' experimental population | -0.482 | 0.618 | 0.526 | -0.917 | 0.359 |
| Male leaving time:Experimental population | -0.206 | 0.814 | 0.251 | -0.823 | 0.411 |
| Male leaving time:Transferred? | 0.286 | 1.295 | 0.299 | 0.864 | 0.387 |

Both males (hazard ratio = 0.198, Wald = 2.625, *P* = 0.009) and females (hazard ratio = 1.492, Wald = 3.244, *P* = 0.001) stayed with their broods for less time when the brood was larger, regardless of their experimental population of origin. Females from block 1 also left significantly earlier than those in block 2 (hazard ratio = 0.576, Wald = -2.166, *P* = 0.030).

We found no evidence that female leaving time was explained by male leaving time. Additionally, the female’s leaving time was not significantly influenced by the interaction between the male’s leaving time and the pair’s experimental population of origin. There was also no evidence that transferring the carcass affected the duration of parental care in Full Care pairs more than No Care pairs: the interaction between whether the carcass was transferred and the pair’s experimental population of origin did not significantly affect the duration of either male or female care (Table 3).

## Discussion

We investigated the extent to which pre-hatching care in carrion nest preparation mediates post-hatching interactions between the family and the nest, and within the family, by focusing on three specific questions.

### Pre-hatching care and offspring performance

How does evolved change in pre-hatching care in the No Care populations combine with post-hatching care to influence offspring performance? We expected that broods derived from each experimental population might generally perform better on No Care prepared carcasses, yielding more numerous and heavier larvae at dispersal. This hypothesis was based on Duarte et al.’s (2021) work on the same experimental populations as we studied, showing that by Generation 14 evolutionary No Care beetles cut an access hole for larvae in the carcass earlier and prepared rounder nests than Full Care beetles. However, by Generation 42, we found no evidence that larvae raised with post-hatching care on these nests attained a greater mass by the time they had completed development. We suggest three explanations. 1) Parents modulate their supply of post-hatching care in relation to the quality of the carcass preparation. In other words, parents simply downgrade their parental care on better carcass nests meaning that there is no additive effect of good carcass preparation and good parental care on larval quality. Our results are only partially compatible with this interpretation: carcass nests prepared by No Care parents were attended for a similar length of time by males, whether males were originally derived from the No Care or the Full Care populations. The nests that induced the greatest duration of post-hatching care were indeed those prepared by Full Care parents. The complicating factor is that this was true only when Full Care males used them for raising larvae. For reasons that are unclear, nests prepared by Full Care parents were attended for far less time when No Care males carried out post-hatching care.

An alternative interpretation is that 2) No Care parents prepare their carcass nests more rapidly but the nest is not ‘better’ in any other sense. Maybe the No Care pairs’ speed of carcass preparation improves larval survival if parents are not present (as in Duarte et al.’s experiments), because when an access hole in inserted before larvae hatch they are better able to establish themselves on a carcass in the absence of parents (Duarte et al., 2021; Eggert et al., 1998). However, if parents are present to assist offspring when they reach the carrion nest then these benefits are no longer discernible. This interpretation is further supported by work from Capodeanu-Nägler et al. (2016) which found that in the presence of post-hatching care there was no difference in brood mass between larvae presented with an unprepared mouse and a prepared carcass nest. This seems like the more plausible of the two explanations to us, based on the evidence that is currently available. Alternatively, 3) since carcass preparation speed and roundness were last assessed at generation 14, perhaps by generation 42 No Care pairs no longer prepared carcasses quicker and more effectively than Full Care pairs, and therefore we see no difference in larval mass because there is no difference in carcass nest quality. However, Duarte et al. (2021) hypothesised that the findings at generation 14 were the result of the selective pressure on No Care pairs to prepare carcasses quickly and effectively to compensate for a lack of post-hatching care, and from generation 14 to 42 this pressure was maintained. Therefore, we have no reason to think that this effect would have been disappeared, but acknowledge that without comparable data we cannot guarantee this to be the case.

#### Are families co-adapted to their carrion nest?

We found experimental evidence consistent with this possibility because when we perturbed this environment by cross-fostering nests between pairs, broods performed less well. However, we found no evidence to suggest that the extent of co-adaptation within each family diverged between the two types of experimental population. Without further experiments, we cannot identify exactly how families are co-adapted to their nests and nest environment.

It might be that any co-adaptation is mediated by chemical cues, that are laid either on the carrion itself or in the surrounding soil, to guide larvae to the carrion nest (Gruszka et al., 2020; Trumbo et al., 2021). Maybe the chemical microenvironment in and around the nest is as vital to the nest’s function as - say - the wall of a bird nest. Perhaps moving nests between boxes was as disruptive as tearing off part of a bird’s nest. However, since we picked up and immediately replaced the carcasses in the broods that did not have their nests translocated, there was the same amount of disturbance across all broods, and thus if it were simply that lower brood mass was caused by disruption of the chemical microenvironment, we would not see a significant effect of translocation. Alternatively, it could be that this chemical environment is more subtly variable among families. When a pair of beetles prepares a carcass they manipulate the microbial volatiles that it releases (Trumbo et al., 2021) – perhaps in ways that are unique to each family. For example, causing a mismatch between the chemicals that the larvae are adapted to and the ones that were present on the carcass in the translocated broods it may have affected larvae by slowing down the rate at which they found the carcass after hatching.

Another suggestion is that parents and larvae are co-adapted to their carcass via the specific community of microbes that parents curate on their carcasses. This could have affected the behaviour of the breeding adults if they were able to detect differences between the microbial community they smeared on the carcass and those on the new carcass they were given, and reduced their parental investment accordingly. This may have evolved as a mechanism for detecting co-breeding, which occurs in *Nicrophorus* species (Komdeur et al., 2013; Müller et al., 1990; Scott, 1997; Sun et al., 2014; Trumbo & Fiore, 1994), and could enable reduced investment in larvae that are not direct descendants (e.g. Richardson and Smiseth (2020), Ma et al (2022)).

More generally, parental investment in birds is known to be affected by whether pairs breed communally or alone, and also by the number of additional breeding pairs. The effects are varied depending on species, with some species increasing investment to compensate for the costs associated with communal breeding, such as in Greater Anis *Crotophaga major* where early-laying females lay larger clutches to compensate for high egg rejection by other females (Riehl, 2010). However, in other species, breeding pairs may reduce their investment, such as in Taiwan Yuhinas *Yuhina brunneiceps* where alpha males and females reduce their incubation effort as the breeding group size increases (Yuan et al., 2005).

Alternatively, it may be that families are co-adapted to the specific cocktail of microbes and proteins in the fluids that their family members deposit onto the carcass and if they receive a carcass prepared by unrelated individuals they do not have optimal developmental resources and suffer slight reductions in growth. There is evidence from other taxa that the interruption of the vertical transfer of microbiota can cause fitness costs in offspring (Greer et al., 2020; Hosokawa et al., 2006; Schwab et al., 2016) but these studies have completely prevented the transmission of either a specific obligative symbiont or all microbes from a particular transmission route. In the case of our study each brood had access to an intact carcass nest microbiome, but potentially not one that was bespokely attuned for their family.

It should also be noted that, although we found evidence of co-adaptation to the nest environment, the effect on brood mass was relatively small (on average less than 0.05 g per brood), indicating that co-adaptation contributes to fitness in a relatively minor way.

#### Does the carrion nest mediate interactions between the parents?

We found that No Care males remained with the brood for the least amount of time. However, No Care females did not adjust their levels of care in response, suggesting they were relatively insensitive to the levels of care supplied by males. Furthermore, the extent of insensitivity to male parental behaviour was not affected by whether their carrion nest was prepared by the focal female or translocated to her. Nor did it differ between No Care and Full Care populations. Both parents instead modulated the duration of their care in relation to brood size, spending less time with broods that were larger.

Although the carrion nest had relatively little effect on interactions between males and females it did appear to mediate interactions between parents and offspring. When parents prepared a larger carcass they produced heavier, though not larger, broods even when the original larger carcass was subsequently transferred away from them and parents were given a smaller carcass nest upon which to raise their offspring instead. Therefore, each individual larva was on average heavier. One explanation is that parents consumed more on the larger carcass themselves whilst they were preparing the nest. If the extent of post-hatching provisioning is state-dependent, then perhaps this means parents were better able to feed offspring after hatching. Alternatively, or as well, parents simply needed to feed less on the carcass themselves after hatching (Boncoraglio & Kilner, 2012; Keppner & Steiger, 2021; Pilakouta et al., 2016) – leaving more resources for their offspring to consume. This is also supported by our result that the longer that males lingered with their offspring, the fewer young they produced (Table 2), which suggests that parents and offspring are in competition for resources on the carcass.

The general picture emerging from these results is that the carrion nest regulates burying beetle family life, but mainly due to traits that are plastically expressed. By mechanisms that remain to be deduced, individual families are co-adapted to their specific nest environment, and/or the immediate environment in the surrounding soil. After hatching, the carrion nest also indirectly influences interactions between parents and offspring, because it is a source of competition between them. However, we found no evidence of divergent evolution in these plastic traits between our experimental populations. Although we have previously shown that No Care populations rapidly evolved to make a nest more quickly for their young (Duarte et al., 2021), here we find that this does not increase larval mass when they do receive post-hatching care and that the role of the nest in family life is seemingly otherwise unchanged. This study represents an advancement in our understanding of the way in which the nest environment mediates family life. Experiments have previously been conducted in birds where nestlings have been reared in a foreign nest, but in those systems the effects of nest construction and parental care are still confounded because the offspring are tended to by the nest builders, rather than their biological parents (e.g. Hõrak et al., 2000; Nilsson & Gårdmark, 2001; Slagsvold & Wiebe, 2007; Teather, 1992). By using the burying beetle as our study system we were able to tease apart these intricately interdependent aspects of parental care and determine to what extent each contributes to correlates of offspring fitness.

## Supporting information

Title page

## Data availability statement

Data have been made available to reviewers and will be made openly available on Dryad at the time of publication.

## Appendix

**Table A1:** Independent variables included in models and the biological reason for their inclusion.

| Independent variable | Reason for inclusion in models |
| --- | --- |
| Current carcass mass | Although we controlled carcass mass experimentally to within 3 g, previous research on *Nicrophorus vespilloides* shows that small differences in carcass mass can lead to significant differences in brood size and mass (e.g. Pascoal et al., 2018; Rebar et al., 2022) |
| Original carcass mass | We included this term to investigate whether the original mass of the carcass could explain any variation in brood size or mass. Previous work has shown that parents cannibalise first instar larvae based on the size of the carcass (e.g. Bartlett, 1988). |
| Parents' experimental population | Central to question 1 in the manuscript |
| Carcass preparers' experimental population | Central to question 1 in the manuscript |
| Carcass transferred? | Central to question 2 in the manuscript |
| Male leaving time | Central to question 3 in the manuscript |
| Female leaving time | Central to question 3 in the manuscript |
| Block | We included this term to test whether any differences we found were due to adaptive evolution (significant population effect) or drift (significant block effect). Since the experimental design had only two blocks, we did not treat block a random effect. |
| Parents' experimental population:Carcass transferred? | This term was included to test whether divergent regimes of post-hatching care caused populations to react differently to being presented with a foreign carcass nest. |
| Original carcass mass:Current carcass mass | This term was included to test whether any effects of original carcass mass were more pronounced when there was a greater difference in mass between the original and current carcass. |
| Parents' experimental population:Male time | Based on previous work (Bladon et al., 2023) we expected that No Care males would leave earlier than Full Care males and so tested for that possibility here. |
| Parents' experimental population:Female time | This term was included to test whether No Care and Full Care females differ in their duration of care. Previous work suggests not (Bladon et al., 2023). |
| Parents' experimental population:Carcass preparers' experimental population | This term was included to test for divergent local adaptation of experimental populations to the type of nest environment they each construct. |
| Current carcass mass:Parents' experimental population | We included this term because previous work suggests that the experimental populations differ in their efficiency of converting carrion into larvae (Schrader, Jarrett, et al., 2015a). |

**Table A2:**
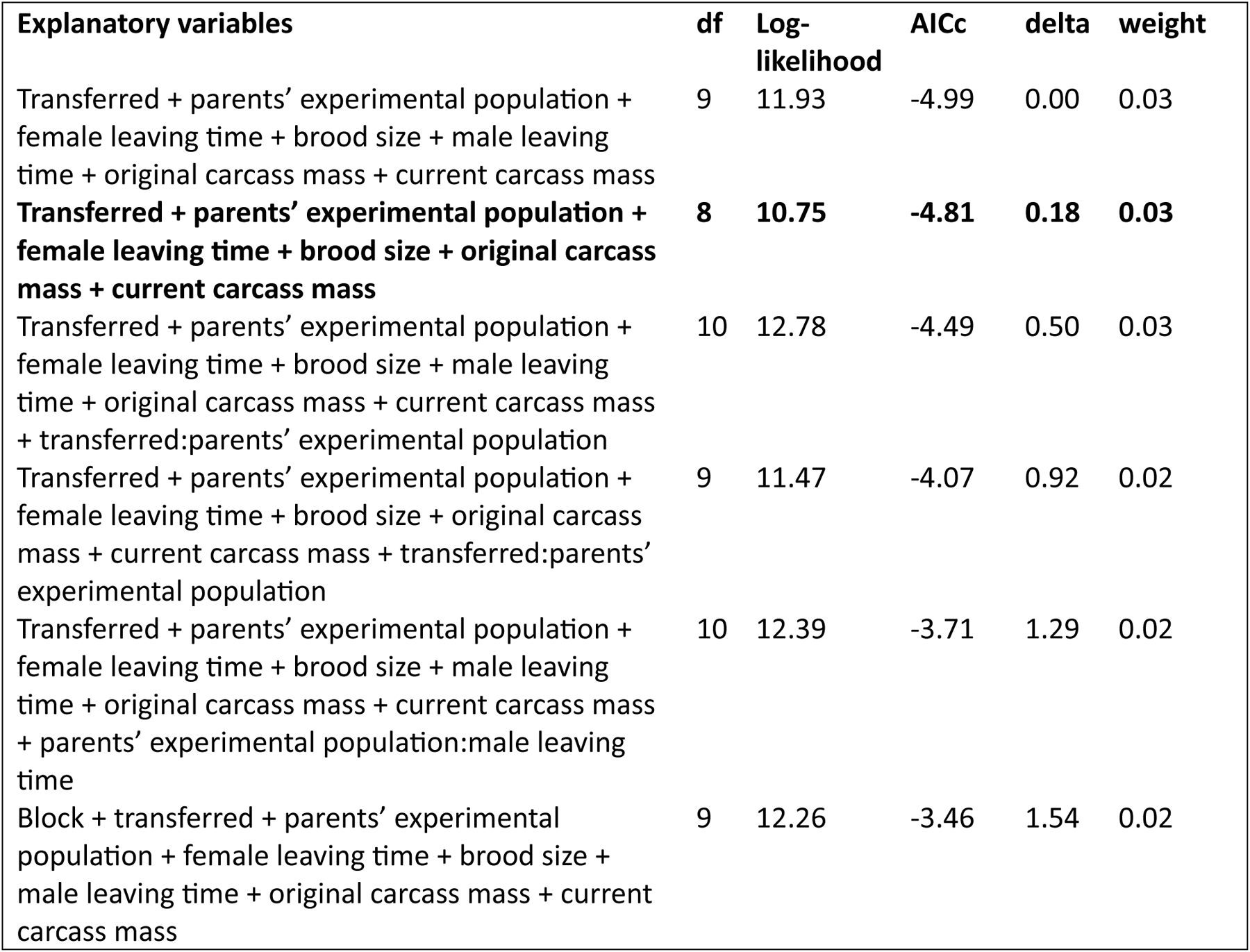
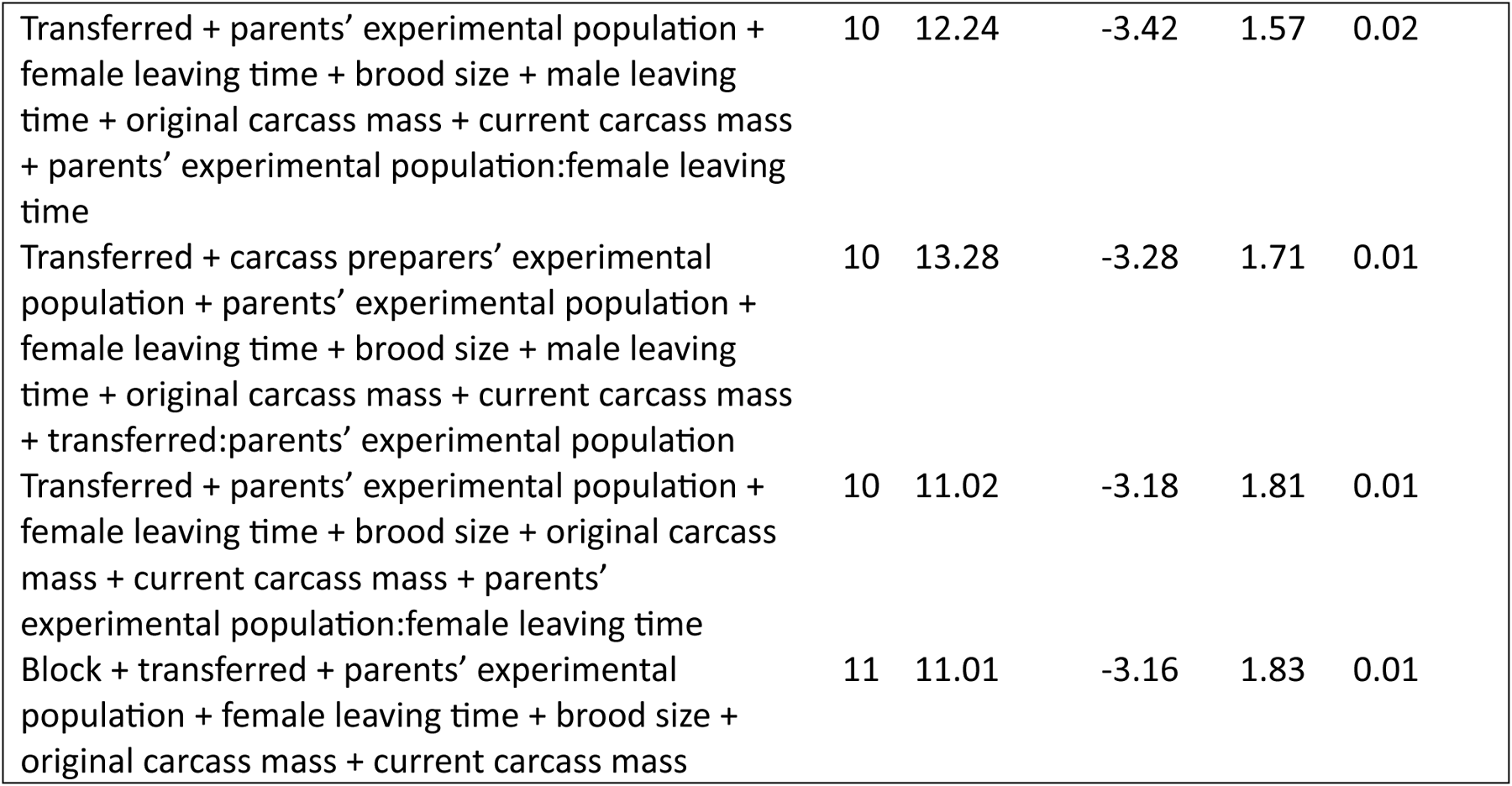
Model selection based on AICc for linear regressions of brood mass at dispersal. Models are ordered by AICc, with only models within two AICc points of the model with the lowest AICc shown. The optimal model is in bold. “:” indicates an interaction.

**Table A3:** Model selection based on AICc for linear regressions of brood size at dispersal. Models are ordered by AICc, with only models within two AICc points of the model with the lowest AICc shown. The optimal model is in bold. “:” indicates an interaction.

| Explanatory variables | df | Log-likelihood | AICc | delta | weight |
| --- | --- | --- | --- | --- | --- |
| Block + female leaving time + male leaving time + current carcass mass | 6 | -684.47 | 1381.34 | 0.00 | 0.05 |
| **Block + male leaving time + current carcass mass** | **5** | **-686.19** | **1382.65** | **1.32** | **0.03** |
| Block + female leaving time + male leaving time + original carcass mass + current carcass mass | 7 | -684.16 | 1382.85 | 1.51 | 0.02 |
| Block + female leaving time + current carcass mass | 5 | -686.32 | 1382.93 | 1.59 | 0.02 |
| Block + transferred + female leaving time + male leaving time + current carcass mass | 7 | -684.23 | 1383.00 | 1.67 | 0.02 |
| Block + parents' experimental population + female leaving time + male leaving time + current carcass mass + parents' experimental population:female leaving time | 8 | -683.25 | 1383.20 | 1.86 | 0.02 |
| Block + carcass preparers' experimental population + female leaving time + male leaving time + current carcass mass | 7 | -684.34 | 1383.22 | 1.89 | 0.02 |
| Block + parents' experimental population + female leaving time + male leaving time + current carcass mass | 7 | -684.37 | 1383.27 | 1.93 | 0.02 |

**Figure A1:**
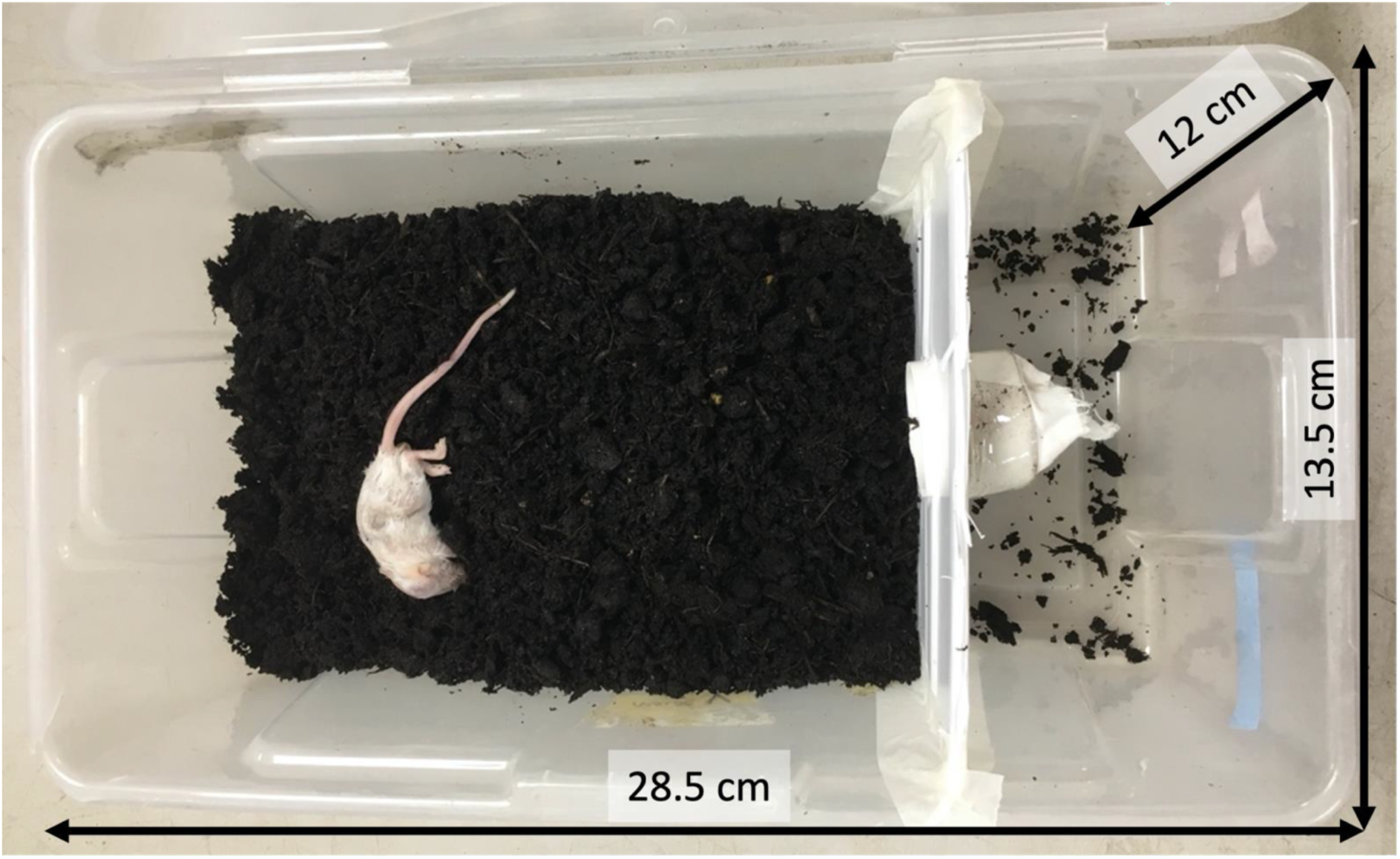
The breeding boxes used in the experiment, containing a breeding compartment (left) and an “escape chamber” (right).

**Figure A2:**
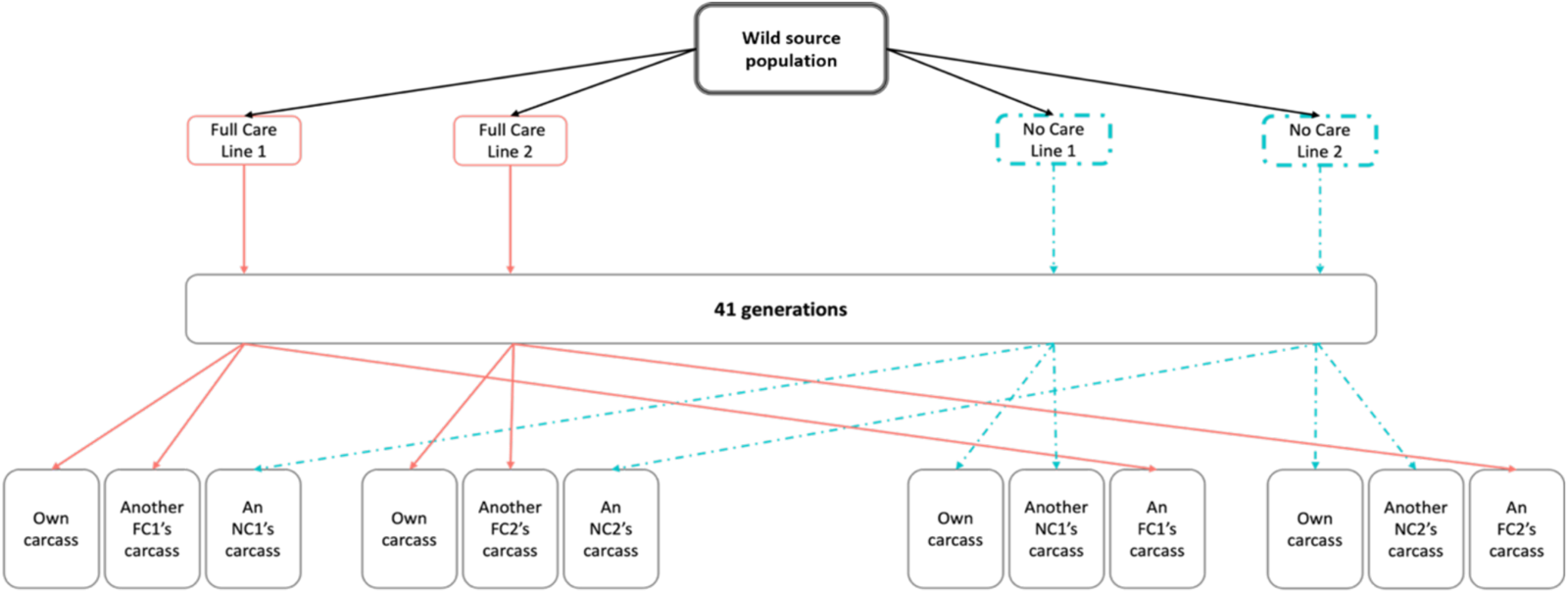
The design of the experimental evolving populations and carcass translocation experimental design.

