## Supplementary material for "Nest construction and its effect on post-hatching family life in the burying beetle *Nicrophorus vespilloides*": Title page


**Author names and affiliations:**

Dr Eleanor K. Bladon[1]* & Professor Rebecca M. Kilner[1]

[1]Department of Zoology, University of Cambridge, Downing Street, Cambridge, CB2 3EJ, UK.


**Word count:** 5,576 (excluding abstract, figure legends, references and supplementary material)


**Acknowledgements:**

This project was supported by a Consolidator's Grant from the European Research Council (310785 Baldwinian_Beetles), by a Wolfson Merit Award from the Royal Society, The Leverhulme Trust (RPG-2018-232), and The Isaac Newton Trust (18.23(q)), each to R.M.K. E.K.B. was supported by a Biotechnology and Biological Sciences Research Council PhD studentship (BB/M011194/1).

We thank Benjamin Jarrett, Darren Rebar, Matthew Schrader and Rahia Mashoodh for establishing and maintaining the experimental evolution lines, and Chris Swannack and Sue Aspinall for beetle maintenance during the initial generations of experimental evolution.
